# Plasticity in evolutionary games

**DOI:** 10.1101/509604

**Authors:** Slimane Dridi

## Abstract

The ability to respond appropriately to environmental cues is fundamental to the success of all forms of life, but previous theoretical studies of the evolution of plasticity make so diverse assumptions that the conditions under which plasticity can emerge in evolving populations are unclear when fitness is frequency-dependent. We study the effect of adding plastic types to symmetric evolutionary games. Since frequency-dependence induces an evolutionary change in the environment of players, one might expect that plastic individuals who can adapt their phenotypes to the environment could have a fitness advantage over simpler purely genetically determined phenotypes. In our model, plastic individuals can detect the type of their opponent before an interaction and condition their action on it. Even though it might appear as an outstanding advantage, such an ability cannot guarantee global stability in all games for even the smallest positive plasticity costs. We identify classes of games where plasticity can or cannot be globally or locally stable. In games where the standard replicator dynamics converge to a pure state, costly plasticity cannot invade an equilibrium population. Costly plasticity can however be locally stable, but the way to achieve this is not to play the best-response to any possible encountered type. Rather, part of the stability success of plastic types is based on establishing Pareto-efficiency as residents; in certain social dilemma games such as the Prisoner’s Dilemma and the Snowdrift game, this implies that plastic types need to cooperate with a strictly positive probability in order to ensure their stability against invasion by pure types. Zero-sum games always allow for the global stability of plastic types. This study offers a more principled way of thinking about the evolutionary emergence of plasticity in social scenarios, and helps demonstrate that such an emergence is strongly dependent on the type of game individuals are faced with.

## 1 Introduction

Most phenotypes are plastic; indeed, in general traits are only expressed in reaction to an environmental cue. The immune system is programmed to detect pathogens, quorum sensing in bacteria conditions gene expression on cell density, plant growth depends on external light via photsynthesis, etc. Being plastic thus seemingly provides a selective advantage to biological organisms. The conditions that favor plasticity evolution have been studied in detail in evolutionary biology and a general conclusion that can be drawn from classical theory is that one requires a varying environment for plastic traits to provide a fitness benefit (Gavrilets and Scheiner, 1993a,*b*; Gomulkiewicz and Kirkpatrick, 1992). Most of these classical models however were developed under the assumption that fitness depends on the environment but not on the phenotype of other organisms in the population, i.e., in the absence of frequency-dependence. It is an open question under what conditions plasticity emerges in frequency-dependent scenarios. In particular, it is unclear how the idea that varying environments favor plasticity evolution can be generalized to a scenario where there is frequency-dependent selection. In order to address this question, one useful way to think about frequency-dependent evolution is to use evolutionary game theory (Taylor and Jonker, 1978; Hofbauer and Sigmund, 1998; Sandholm, 2011). Interactions between individuals can be described as games where every individual has a set of strategies, and the fitness of individuals involved in the game is dependent on everyone’s strategy. In this context, the idea of a varying environment can take several forms. A first one is that the rules of the game change over evolutionary time, possibly under the influence of the population’s behavior itself, which can occur typically under competition for limited resources (Dridi and Lehmann, 2014; Weitz et al., 2016; Hilbe et al., 2018). A second and simpler form of environmental variation in social evolution however is to notice that frequency-dependence itself induces environmental variation for a focal individual or type (Graves and Weinreich, 2017). Indeed, in frequency-dependent selection, the frequency of other types makes part of the environment, and since the frequencies evolve over time, the environment of any focal individual can be seen as varying between generations. Moreover, in models that explicitly use normal-form games to capture social interactions, one generally needs to make assumptions on how population members are matched to interact in a game. Very often, random matching between players is assumed, which also creates within-generation fluctuations in the environment of a focal player because a player is generally assumed to not know who is going to be its sampled opponent. These observations suggest that even in the absence of changes in the structure of the evolutionary game determining individuals’ fitness, there is still environmental variation and potentially a selection pressure in favor of plasticity. In order to further examine this question, one must determine what it means to have a plastic phenotype in a game theoretical context.

Plasticity in social evolution has been studied to address many different questions, from studies of co-operation in the iterated prisoner’s dilemma to investigations of signalling, and learning (Axelrod and Hamilton, 1981; McElreath and Boyd, 2007; Stewart and Plotkin, 2013; Dridi and Lehmann, 2015, 2016). In studies of the iterated prisoner’s dilemma, focus has been given to repeated game strategies, and in particular to memory-one automata that condition their action at the current round on the outcome of the game at the previous round. Well-known strategies such as Tit-for-tat, Win-Stay-Lose-Shift, Zero-Determinant strategies all belong to this class. In studies of the evolution of signalling, one generally tries to understand the principles leading to the evolution of the ability to manipulate what others detect from one’s own type. This goes well beyond our mere interest in the emergence of plasticity, because for such manipulation to take place, one must assume the pre-existence of plasticity in social strategies. Other researchers have focused on learning strategies, where the entire history of the game determines the current action of a player. This focus stems from the observation that reinforcement learning underlies virtually all animal behaviors. These studies have provided interesting insights into our understanding of how natural selection shapes complex strategies for repeated interactions. However, complex strategies that allow an individual to condition behavior on the environment, on opponents’ behavior, or on memory of past events are only possible if, in the first place, individuals possess the ability to express plastic social phenotypes. Previous work seemingly does not address the question of the evolutionary emergence of social plasticity, so it remains unclear what is the main advantage of plasticity in social evolution.

If the success of plasticity should depend on the exact way it is implemented, these different implementations can possibly classified according to the complexity of the resulting strategy (e.g., always fleeing when faced with a predator is probably not as complex as learning to hide food to prevent conspecifics from stealing it; Emery and Clayton, 2001, 2004, 2009). In order to investigate the evolutionary emergence of plasticity, one must consider its simplest possible form. In this paper, we will adopt one of the simplest implementations of plasticity that we can think of, namely we will assume that plastic individuals can detect the type of their opponent before an interaction takes place, and can condition their action on the detected type. At first, this might seem as a considerable advantage, but we will see below that even the smallest cost impedes plastic types to dominate in all circumstances. Being able to detect others’ types is an assumption that can be justified on several grounds. If the evolving behavioral trait of interest is linked to morphological or hormonal characteristics, then it could be possible for a plastic type to base their strategy on, e.g., vision or smell. If outcomes of games are public information and the game is repeated (but not necessarily with the same partner) during individuals’ lifespan, then our model can be viewed as a generalisation of studies of the evolution of cooperation through reputation (Nowak and Sigmund, 1998). Yet another possibility is that the game is repeated between the same players, and our implementation of plasticity might correspond to a learning strategy that consists in uncovering the pure strategy of the opponent, and then playing an appropriate response (Dridi and Lehmann, 2014). However, in practice when analysing our model we abstract away from details about how the plastic type reaches its behavioral response.

Even though we try to adopt a simple form of plasticity and do not focus on the mechanistic details leading to plasticity, one potential concern that may arise at this point is that the genetic and molecular machinery necessary to perform a combination of strategy detection and appropriate response might be very complex to evolve, even for the most basic forms of plasticity. A perfect response to existing types is unlikely to emerge out of a background of pure genetic determination and previous research indeed suggests that such perfect responses might be very difficult to evolve (McNamara et al., 1999; André and Day, 2007). For this reason, we will allow our plastic types to adopt any possible response to their opponents, in contrast to a previous work on the topic (Banerjee and Weibull, 1995), and we will study how the evolutionary success of plastic types depends on their response to pure types. Moreover, the perceptual system allowing one to infer others’ strategies might at first also be defective if it evolves from a state where there was no perceptual system of this kind in the ancestral population. We capture such imperfections by imposing a fitness cost on the expression of the plastic phenotype.

In the rest of the paper, we define a model that makes our assumptions more precise, and analyze the evolutionary performance of plastic types when pitted against individuals who can only express a fixed pure strategy in a normal-form game. We pay particular attention to scenarios where plasticity can invade a population of pure types and eliminate them such that it becomes the evolutionarily stable strategy. We also give a special focus to 2 *×* 2 games and analyze the replicator dynamics for four standard games of cooperation: the Prisoner’s dilemma, the Stag-hunt game, the Snowdrift game, and a Mutualism game. We also provide classes of games where plastic types can or cannot be globally or locally stable under the standard replicator dynamics.

## 2 Model

We consider a standard model of a well-mixed population in which players are matched randomly in pairs to play a 2-player *n*-action game. We denote the set of actions by 𝒜 with |𝒜| = *n*. The population consists of *n* + 1 types: the first *n* types correspond to each pure action while the (*n* + 1)-th type can detect others’ type before choosing an action. This plastic type, denoted *p*, has a strategy described by the vector **z** = (**z**_1_,…, **z**_*n*_, **z**_*n*+1_), where **z**_*i*_ = (*z*_*i*1_,…, *z*_*in*_) ∈ Δ𝒜 is the mixed strategy adopted by *p* when faced with type *i* ∈ *𝒜 ∪* {*p*}. The symbol Δ 𝒜 denotes the *n*-dimensional simplex, such that *z*_*ik*_ is the probability that the plastic type plays strategy *k* against type *i*. With these definitions, the cartesian product 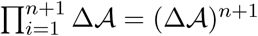 is the strategy set of the plastic type. The mixed strategy **z**_*n*+1_ = **z**_*p*_ ∈ Δ*A* is the strategy adopted when faced with another plastic individual. The payoff of type *i* ∈ 𝒜 against type *j* ∈ 𝒜 is denoted *π*(*i, j*). For interactions involving the plastic type *p*, we mostly write the payoff *π*(*p, i*) = *π*(**z**_*i*_, *i*) to emphasize the dependence of the payoff on the mixed strategy **z**_*i*_ of type *p* against *i*. We identify the payoffs *π*(*p, i*) and *π*(*i, p*) with the expected payoff generated by the mixed strategy of the plastic type, that is

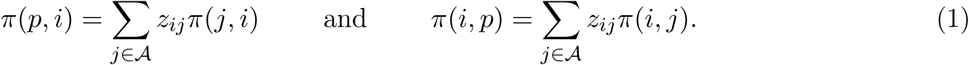

When two plastic individuals meet, their payoff reads

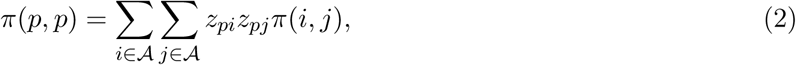

where we assumed that both plastic types adopt the same responsive strategy. We do not consider selection on the plastic response itself, **z**, in this paper, so all plastic types will always have the same responsive strategy. We are interested in tracking the vector of frequencies of the types **x** = (*x*_1_,…, *x*_*n*_, *x*_*n*+1_) ∈ Δ^*n*+1^ such that 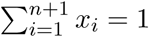. The frequencies evolve according to the standard replicator dynamics. We write *w*_*i*_(**x**), *i* = 1,…, *n*, for the fitness of type *i* when the population is in state **x** which is calculated as the average payoff at state **x**, or

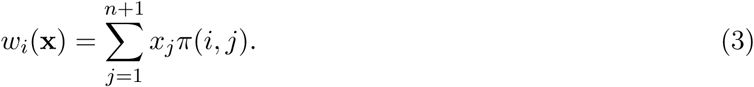

We further assume that type *p* pays a cost *k >* 0 for expressing a plastic response so that its fitness reads

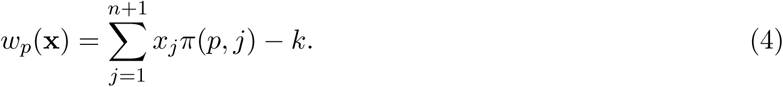

The frequency of any type *i* evolves according to the replicator dynamics, which are given by the differential equations

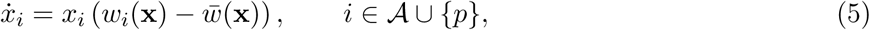

where

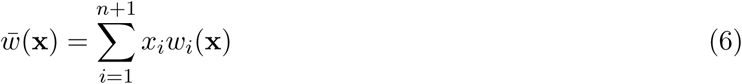

is the average fitness in the population at state **x**. A further element of notation is that we denote by *ϕ*(*·*, **x**_0_): ℝ → Δ^*n*+1^ the global solution trajectory to eq. 5 that passes through **x**_0_, which means that *ϕ*(*t*, **x**_0_) is the vector of frequencies at time *t* given that the vector frequencies passes through **x**_0_. We denote by *ϕ*(*t*, **x**_0_)_*i*_ the *i*^th^ element of *ϕ*(*t*, **x**_0_), with *i* ∈ *𝒜 ∪* {*p*}.

## 3 Optimizing the success of plasticity

Consider the following evolutionary scenario. The population consists of pure types only and is at equilibrium of the replicator dynamics (eq. 5). A mutant plastic type with arbitrary response **z** is introduced in the population. Will the mutant frequency initially grow? Can this mutant plastic type reach fixation and eliminate pure types? Clearly, the answer to these questions will depend on the strategy, **z**, of the plastic type. A major goal of this paper is to find the conditions under which global stability of plasticity is possible. Namely we are interested in the conditions for which lim_*t*→*∞*_ *ϕ*(*t*, **x**_0_) = **e**_*p*_, for all **x**_0_ ∈ int(Δ^*n*+1^). In this section, we distinguish two different concepts, which will prove useful in the following analyses aimed at the aforementioned goal: (1) the best outcome that plasticity can achieve in any game; (2) the minimum strategy required for plasticity to be globally stable in a given game. We will see in our results below that in some games the best outcome that plasticity can achieve is not enough to survive natural selection. In other games, plastic types do not need to do their “best” in order to achieve global stability. Below, we define more precisely what we mean by the plastic types doing their best. But we already note at this point that the problem of finding a plastic strategy that grants global stability can be divided into two different tasks. The first one is to maximize the chances of a plastic type invading all equilibrium populations of the pure type game, and the second one is to find a strategy that guarantees the stability of the plastic type against pure invaders.

### 3.1 Optimizing invasion success

It seems obvious that in order to maximize its invasion success of monomorphic populations, a plastic type must choose the best-response to each existing type, i.e., for each *i* ∈ 𝒜 the plastic type should choose 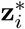 such that

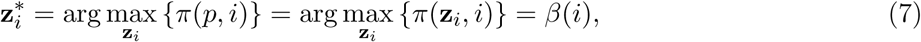

where we used *β*(*·*) to denote the best-response function. If the strategy indeed maximizes the chances of invasion of the plastic type, it does not guarantee invasion. Also, it is not necessary to play 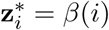 in order to invade a monomorphic population of a pure type. In general, the condition for invasion is given by solving the inequality

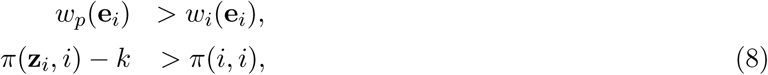

where the *i*-th basis vector **e**_*i*_ ∈ Δ^*n*+1^ is the state where the population consists only of type *i*, i.e., such that *x*_*i*_ = 1 while *x*_*j*_ = 0, *i ≠ j*. The above condition can only be true when strategy *i* is not a best-response to itself, i.e., when (*i, i*) is not a pure symmetric Nash Equilibrium (NE). Hence Proposition 2 below. For a population at a mixed equilibrium, **x**_NE_, let *π*(*i*, **x**_NE_) = *π*(**x**_NE_) = *w*(**x**_NE_) be the fitness or expected payoff obtained by a pure type *i* in the support of **x**_NE_. Then invasion is equivalent to requiring that

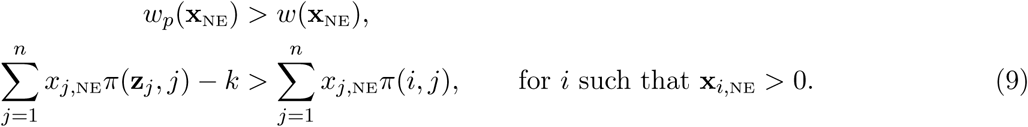

We show in Prop. 3 that there always exists a strategy of the plastic type that allows it to invade the mixed NE. In particular, playing the best-response to every pure type in support of a mixed NE allows plasticity to invade this mixed NE. However, and once again, it is not necessary to play the best-response to pure types in order to achieve this outcome.

### 3.2 Optimizing stability

To optimize its stability, the plastic type must choose a strategy 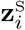 that minimizes the payoff of any pure mutant type when facing the plastic type or

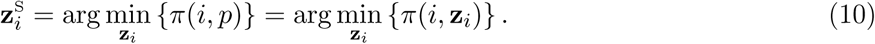

Note that 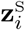 is different than the best-response because it is focused on minimizing the success of mutants. To further optimize its stability the plastic type must choose a strategy against itself 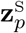 such that

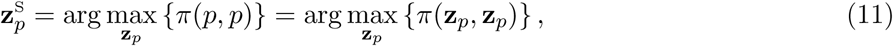

which corresponds to the symmetric Pareto-efficient equilibrium of the underlying symmetric game. We remark that maximizing invasion success might conflict with maximizing stability, i.e., in general **z**^*\**^ *≠* **z**^S^ (we will see that this is so in a Stag-hunt game, for example). Hence, there is no general recipe for the plastic type to achieve global stability in all games. Moreover, even if **z**^*\**^ = **z**^S^, this does not mean that the plastic type will achieve global stability (we will see that this is so in a Prisoner’s dilemma game, for example). It is not necessary to play 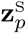, to ensure local stability of plasticity. The condition for stability against a pure type is

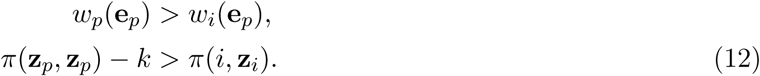

We show in Prop. 4 that eq. 12 implies that a necessary condition for the impossible local stability of plasticity is that (*i, i*) is the symmetric Pareto-efficient outcome. This is a necessary but not a sufficient condition for the instability of plasticity, as we will illustrate when we discuss social dilemma games. Indeed in social dilemma games, plasticity can always be locally stable.

## 4 Conditions for global stability of plasticity

The various observations that we made above put us in the position to state the following result.

### Proposition 1

(Sufficient conditions for global stability of plasticity). *If* ∃**z** ∈ (Δ 𝒜)^*n*+1^ *such that:*

1. *π*(*j, j*) *< π*(**z**_*j*_, *j*), *∀j* ∈ 𝒜
2. *π*(*j*, **z**_*j*_) *< π*(**z**_*p*_, **z**_*p*_), *∀j* ∈ 𝒜
3. 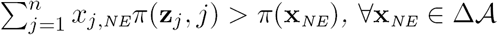

*then there exists a* 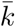 *such that, for all* 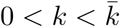, lim_*t*→*∞*_ *ϕ*(*t*, **x**_0_) = **e**_*p*_, *for all* **x**_0_ ∈ int(Δ^*n*+1^),

*Proof.* The set of equilibrium points of the replicator dynamics is the union of the **e**_*i*_, *i* ∈ 𝒜 and of the NE of the game. Adding a plastic type to a game is equivalent to adding an action to the underlying symmetric game, and thus leaves unchanged the original equilibria of the pure-type game, and adds one or more equilibria. In particular, it adds at least one equilibrium, namely **e**_*p*_, the equilibrium consisting of the plastic type only. To show the global stability of plasticity in the game consisting of the *n* types and the plastic type, it thus suffices to show that all the original equilibria of the replicator dynamics are locally unstable, and that the equilibrium **e**_*p*_ is locally stable.

Condition (1) guarantees the instability of all **e**_*i*_, *i* ∈ 𝒜. Indeed, condition 1 implies that there exists a *k*_1_ such that, for all 0 *< k < k*_1_, we have *w*_*p*_(**e**_*i*_) *> w*_*i*_(**e**_*i*_). This inequality in the fitnesses at a monomorphic population of a pure type implies that the equilibrium **e**_*i*_ is unstable (Hofbauer and Sigmund, 1998).

Condition (2) guarantees the local stability of **e**_*p*_ for the same reason that condition 1 guarantees the instability of **e**_*i*_, i.e., there exists a *k*_2_ such that, for all 0 *< k < k*_2_, we have *w*_*p*_(**e**_*p*_) *> w*_*i*_(**e**_*p*_).

Condition (3) finally guarantees local instability of any **x**_NE_ ∈ Δ 𝒜, for the same reason that condition 1 guarantees the instability of **e**_*i*_, i.e., there exists a *k*_3_ such that, for all 0 *< k < k*_3_, we have *w*_*p*_(**x**_NE_) *> w*_*i*_(**x**_NE_) for any *i* in the support of the NE. Taking 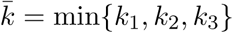 implies that conditions 1, 2, and 3 will be satisfied at the same time for any 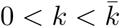, which completes the proof.

Condition 1 of Prop.1 requests that the game possesses no pure symmetric NE, while condition 2 requests that plastic types can play a secret-handshake strategy when meeting themselves that would overcome the possible payoffs of pure invaders. Condition 3 finally requests from plastic types that they play a strategy that guarantees invasion of mixed NE. While we have noted above that playing the best-response to every pure type allows plasticity to invade a mixed NE, we will see with examples that invading a mixed NE is not equivalent to invading all monomorphic populations consisting of strategies in the support of the NE.

## 5 2*×*2 games of cooperation

Since the most famous plastic strategies in evolutionary biology are strategies in repeated games that can sustain cooperation, one is tempted to generalize the idea and ask: Does plasticity generally favor the evolution of cooperation? Or said differently, do plastic types need to establish cooperation in order to successfully invade populations of pure types? We consider in this section four 2 *×* 2 games of cooperation traditionally studied in evolutionary biology: the Prisoner’s Dilemma game (PD), the Stag-hunt game (SH), the Snowdrift game (SD), and the Mutualism game (MG). We label the two possible actions as *C* (for Cooperation) and *D* (for Defection), and this generates a plastic type with a three-dimensional trait *z* = (*z*_*C*_, *z*_*D*_, *z*_*p*_) ∈ [0, 1]^3^, where *z*_*i*_ is the probability to cooperate when facing type *i* ∈ {*C, D, p*}.

We perform a sensitivity analysis with respect to the parameters *z*_*C*_, *z*_*D*_, *z*_*p*_. For each value of these parameters, we determine whether and when plasticity is able to invade equilibrium populations consisting of pure types and/or can resist invasion by pure types. We will use the standard notation for the payoffs of 2 *×* 2 social dilemma games, often used in the biological literature, i.e., *π*(*C, C*) = *𝓡, π*(*C, D*) = *𝒮, π*(*D, C*) = *𝒯*, and *π*(*D, D*) = *𝒫*.

### 5.1 Prisoner’s dilemma game

The payoffs of the Prisoner’s dilemma satisfy 𝒯 > 𝓡 > 𝒫 > 𝒮 as well as 𝓡 > (𝒯 + 𝒮)/2. In this game, there is one dominant action, *D*. The outcome (*D, D*) is the only stable equilibrium of the pure-type game. From condition 1 of Prop. 1, we thus have that plastic types cannot invade a population of defectors, which means that the inequality

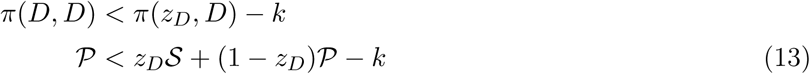

can never be satisfied. But can plasticity invade cooperators, and is it immune against invasion by pure types? As to the first question, if plastic types play *z*_*C*_ = 0 (always defect) against cooperators, they will obviously invade a population of cooperators. But any *z*_*C*_ that satisfies

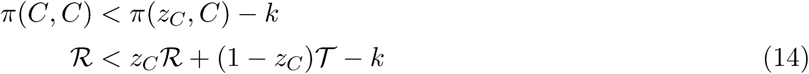

would guarantee that plastic mutants invade a monomorphic population of cooperators. As to the second question regarding the ability of plastic types to resist invasion by pure types, we have to set *z*_*C*_, *z*_*D*_, and *z*_*p*_ so that condition 2 of Prop. 1 holds. In other words we first have to solve the inequality

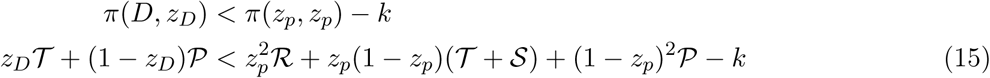

for *z*_*D*_ and *z*_*p*_. In Fig. 1, we show the regions of the space defined by combinations of (*z*_*D*_, *z*_*p*_) that satisfy eq. 15, which constitute the set of plastic types that are immune against the invasion by defectors in the Prisoner’s dilemma game. Note that eq. 15 implies that the plastic type can be immune against the invasion by defectors only if *z*_*p*_ *>* 0 (this can be shown by setting *z*_*p*_ = 0 in eq. 15, which then cannot be satisfied because 𝒯 *>* 𝒫). To establish the local stability of plasticity against all pure types, we also have to solve the same type of inequality involving *z*_*C*_, i.e.,

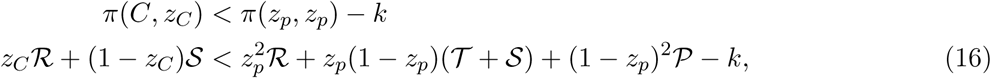

which defines the set of plastic types that are immune against the invasion by cooperators in the Prisoner’s dilemma game.

**Fig. 1:**
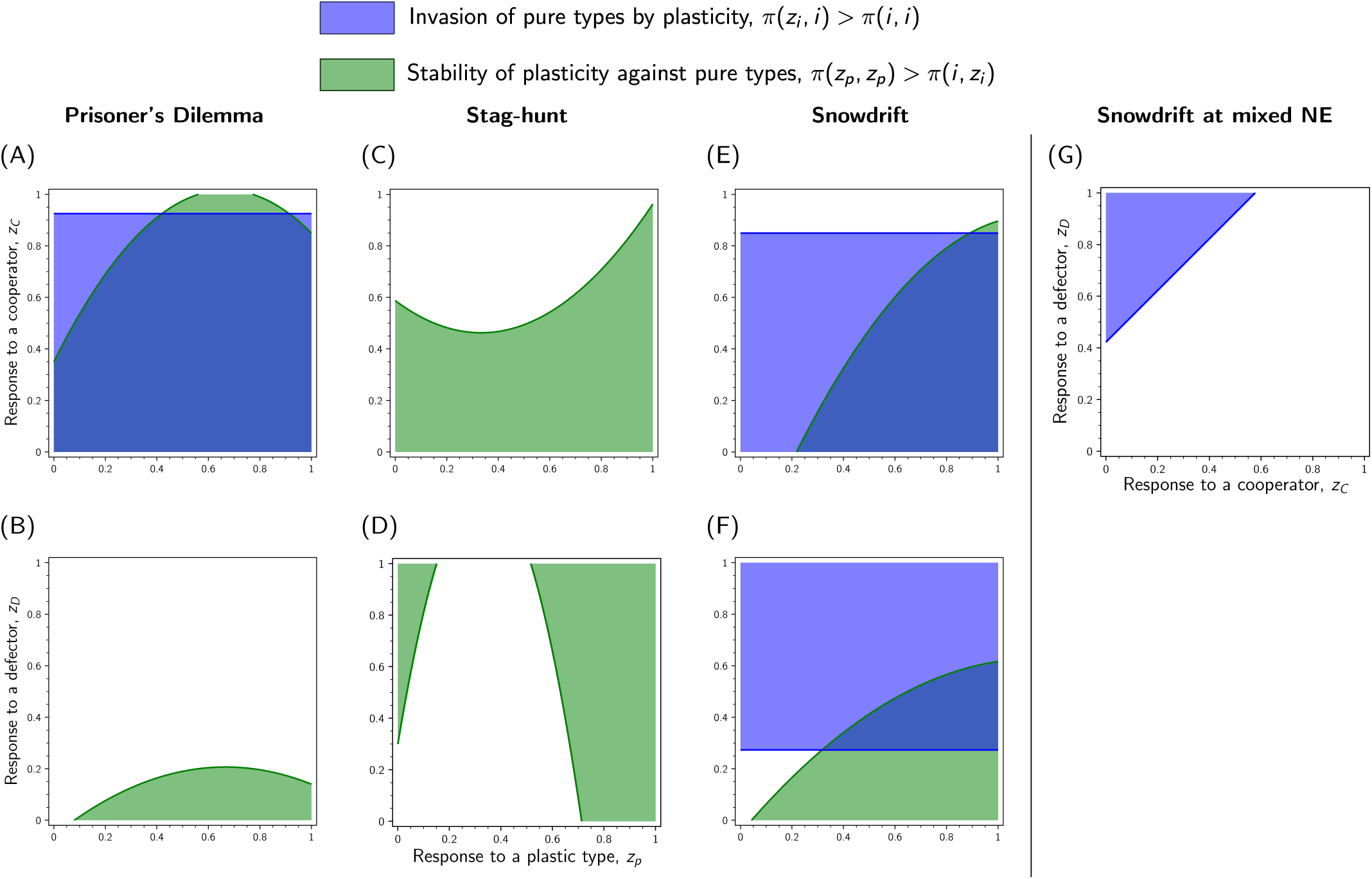
Regions of stability and invasion of plastic types (as defined by eqs. 15–16). (A)–(F) For each social dilemma game (columns) we show the response of the plastic type, that is the combinations (*z*_*p*_, *z*_*C*_) (top row) and (*z*_*p*_, *z*_*D*_) (bottom row) that allow invasion of monomorphic populations of pure types by plasticity (blue), and the ones that allow stability of plasticity against the invasion by pure types (green). We recall here that *z*_*i*_ is the probability to cooperate of the plastic type against each type *i* ∈ {*C, D, p*}. (G) Combinations of (*z*_*C*_, *z*_*D*_) that allow plasticity to invade a population of pure types at the mixed ESS of the Snowdrift game. If (*z*_*C*_, *z*_*D*_) are chosen in the blue region of (G) and (*z*_*C*_, *z*_*D*_, *z*_*p*_) are further chosen at intersection of the green and blue regions of panels (E)–(F), plasticity is globally stable in the Snowdrift game. Parameter values: Prisoner’s dilemma game: (*𝓡, 𝒮, 𝒯, 𝒫*) = (3, 1, 7, 2); Stag-hunt game: (*𝓡, 𝒮, 𝒯, 𝒫*) = (8, 0, 4, 5); Snowdrift game: (*𝓡, 𝒮, 𝒯, 𝒫*) = (3, 0.1, 5, −1).

What are the conditions on the payoffs of the game, (𝓡, 𝒮, 𝒯 𝒫), that allow these inequalities to be satisfied? In Fig. 2, we illustrate the fact that the feasible payoffs of a plastic type as a resident (the right-hand side of eqs. 15–16) are on the line *x* = *y* within the convex hull of feasible payoffs. This means that the payoff *π*(*z*_*p*_, *z*_*p*_) satisfies

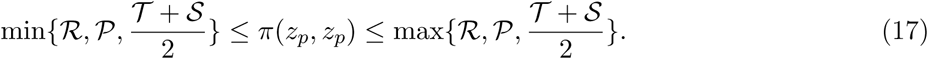

**Fig. 2:**
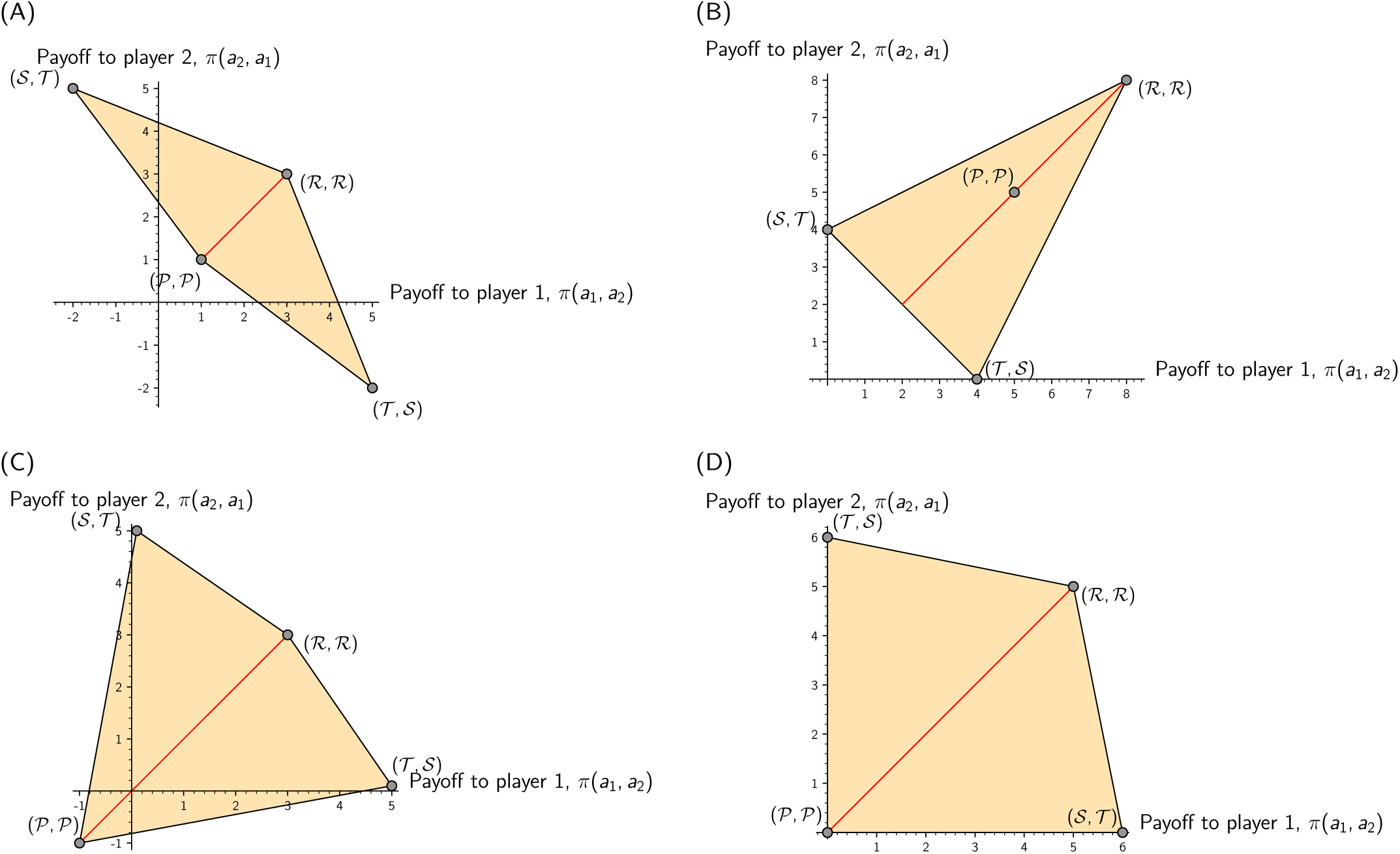
The convex hull of feasible payoffs (orange region) in games of cooperation, and the line (red) of feasible payoffs in a monomorphic population of plastic types. (A) Prisoner’s Dilemma game with (*𝓡, 𝒮, 𝒯, 𝒫*) = (3, −2, 5, 1). (B) Stag-hunt game with (*𝓡, 𝒮, 𝒯, 𝒫*) = (8, 0, 4, 5). (C) Snowdrift game with (*𝓡, 𝒮, 𝒯, 𝒫*) = (3, 0.1, 5, −1). (D). Mutualism game with (*𝓡, 𝒮, 𝒯, 𝒫*) = (5, 6, 0, 0).

Combining the above inequality with eq. 16, we then see that plasticity cannot be locally stable against mutant cooperators if

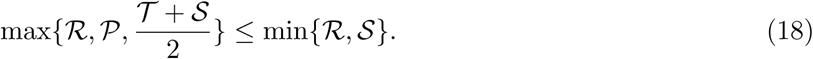

Similarly, using eq. 15, plasticity cannot be locally stable against mutant defectors if

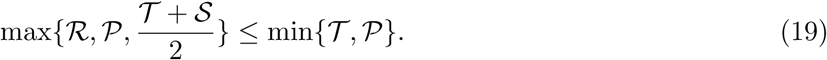

The two latter inequalities are not satisfied in the Prisoner’s dilemma, which means that plasticity can always find a strategy that makes it stable against the invasion by pure types.

### 5.2 Stag-hunt game

We define this game as one with payoffs satisfying 𝓡 > 𝒫 > 𝒯 > 𝒮 and 𝓡 + 𝒮 > 𝒯 + 𝒫, which entails that (*C, C*) and (*D, D*) are two pure NE of the game, with (*C, C*) being the Pareto-dominant equilibrium, and (*D, D*) being the risk-dominant equilibrium. Consequently (*C, C*) and (*D, D*) are both asymptotically stable equilibria of the replicator dynamics in the pure-type game. From condition 1 of Prop. 1, plastic types cannot invade any monomorphic population of pure types. However, solving inequalities 15 and 16, we find that plasticity can be locally stable (Fig. 1). This game is a good illustration of the idea that invasion ability and local stability of our plastic types might be impossible to reconcile. Indeed, the plastic type needs to anti-coordinate with *C* and *D* in order to ensure local stability, but the best it can do in monomorphic populations of either *C* or *D* is to coordinate; and this best effort is not even enough to be able to invade monomorphic populations.

We finally mention a result in the Stag-hunt game that may be of interest beyond evolutionary biology. Namely, in our model, costless plasticity in the Stag-hunt game is a solution to the problem of converging to the payoff-dominant equilibrium. Even though our main focus in this paper is on the case where the cost of plasticity is positive (*k >* 0), we mention this result because most evolutionary or learning processes known to us converge to the risk-dominant equilibrium, (*D, D*) (or stochastic evolutionary processes admit a stationary distribution that puts more mass on the risk-dominant equilibrium). Here, if one sets *k* = 0 and (*z*_*C*_, *z*_*D*_, *z*_*p*_) = (1, 0, 1), then the face of the simplex such that defectors are at frequency 0 is globally stable. All points on this face are neutrally stable, i.e., any mix of cooperators and plasticity is globally stable. In such populations, everyone cooperates and thus achieves the maximum possible payoff in the Stag-hunt game.

### 5.3 Snowdrift game

In the Snowdrift game, the payoffs are such that 𝒯 > 𝓡 > 𝒮 > 𝒫. This game thus calls for closer attention because it has a symmetric mixed NE. In order to achieve global stability, a plastic type must be able to invade any mixture of the pure types, including the NE mixture. This game is in fact an excellent illustration of the idea that playing the best-response to every pure type does not always yield the best outcome for a plastic type. Indeed, in this game, playing the best-response to cooperators and defectors guarantees that plasticity will invade monomorphic populations and the mixed NE, but this strategy does not guarantee local stability of plasticity. However there exists a strategy, which is not the best-response, that guarantees global stability of plasticity, provided the payoffs allow it. Such a strategy is found by solving simultaneously the inequalities in eqs. 13–16, together with the following inequality

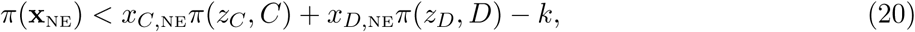

which is the condition for plasticity to invade a population at the mixed NE of the Snowdrift game. It turns out to be quite difficult to reduce the inequalities in eqs. 13–16 and eq. 20, but we provide in Fig. 1 an example showing that there exists a strategy and payoffs of the Snowdrift game such that they all hold, i.e., such that plasticity is globally stable.

The Snowdrift game also provides interesting counter-examples to conjectures that one might be tempted to make: if a plastic type can invade every monomorphic population, it cannot necessarily invade a population at the mixed NE; in Fig. 1, take for example (*z*_*C*_, *z*_*D*_, *z*_*p*_) = (0.2, 0.4, *z*_*p*_). The converse is also false: if a plastic type can invade the mixed NE, this does not mean that it can invade every monomorphic population in the support of the NE. However, in any game with no pure NE, there always exists a strategy that guarantees invasion of both mixed NE and pure monomorphic populations; in particular, playing the best-response against every pure type always guarantees such an outcome.

Another statement that can be proved wrong using the Snowdrift game is that games without pure symmetric NE always make possible global stability of plasticity. In the Snowdrift game, setting *𝒯* very large creates a game that has still no pure symmetric NE, but where there is no longer a strategy that makes plasticity globally stable. The intuitive reason is that setting *𝒯* very large reduces substantially the upper bound on *z*_*D*_ such that plasticity is locally stable, i.e., the (green) region of local stability of plasticity in Fig. 1 shrinks and then no longer intersects the (blue) region where plasticity can invade a monomorphic population of defectors.

### 5.4 Mutualism game

In this game we assume simply that 𝓡 > 0 and 𝒮 > 0, while 𝒯 = 𝒫 = 0, such that cooperation is a dominant action and the only NE of the pure-type game. Despite not being a social dilemma in the sense that there is no conflict between cooperation and rational behavior, this game is still interesting when considering the possible evolution of plasticity. Indeed in this game there are values of the payoffs for which plasticity is neither locally stable nor able to invade a monomorphic population of cooperators, for any value of (*z*_*C*_, *z*_*D*_, *z*_*p*_), which implies that any share of plastic mutants – irrespective of their strategy – will be unable to survive natural selection. From our previous analysis, we know that plasticity cannot invade a monomorphic population of cooperators, since cooperation is a dominant action. Plasticity can however invade a population of defectors by playing a sufficiently high *z*_*D*_. However, if 𝒮 ≥𝓡, then eq. 18 is satisfied, which means that plasticity cannot be locally stable. If, on the other hand, 𝒮 < 𝓡 then a plastic type can resist invasion by cooperators through defection against them. In Fig. 3, we show the phase portrait of trajectories in a particular Mutualism game where 𝒮 ≥𝓡, but also for all games of cooperation studied in this section.

**Fig. 3:**
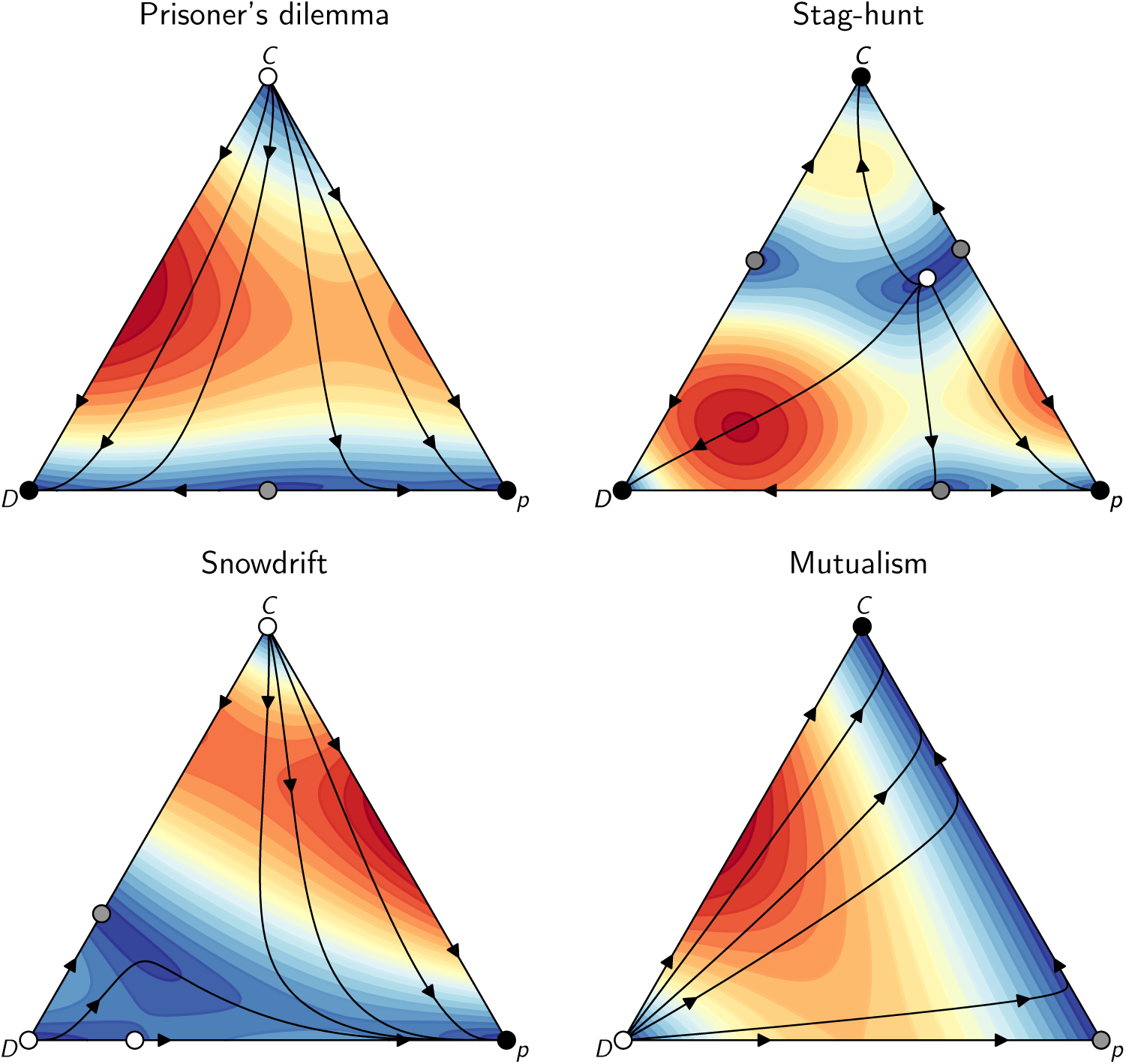
Replicator dynamics with added plastic types in 4 games of cooperation. Each triangle represents a projection of the 3-simplex on the 2-dimensional plane, where at each vertex the corresponding strategy (*C*, *D*, or *p*) is at frequency 1. On the face opposed to a vertex, the corresponding strategy is absent, i.e., it is at frequency 0. These plots are displayed for the best possible outcome that can be achieved by a plastic type. Parameter values: Prisoner’s Dilemma: (𝓡, 𝒮, 𝒯, 𝒫) = (3, −2, 5, 1), and (*z*_*C*_, *z*_*D*_, *z*_*p*_) = (0, 0, 1); Stag-hunt: (𝓡, 𝒮, 𝒯, 𝒫) = (8, 0, 4, 5), and (*z*_*C*_, *z*_*D*_, *z*_*p*_) = (0, 1, 1); Snowdrift: (𝓡, 𝒮, 𝒯, 𝒫) = (2.5, 0.1, 5, −1), and (*z*_*C*_, *z*_*D*_, *z*_*p*_) = (0.03, 0.45, 0.9) [this strategy is chosen in the regions of invasion and stability of Fig. 1]; Mutualism: (*𝓡, 𝒮, 𝒯*, *𝒫*) = (5, 6, 0, 0), and (*z*_*C*_, *z*_*D*_, *z*_*p*_) = (1, 1, 1); in all games we used a cost of *k* = 0.3. Contours indicate speed with blue corresponding to slower dynamics and red corresponding to faster dynamics. Black dot: Asymptotically stable equilibrium (a “sink”) of the replicator dynamics; gray dot: an unstable saddle; white dot: an unstable source.

## 6 More general games

We now state results that hold for more general classes of games than 2 *×* 2 games of cooperation. In all of our results, we assume that the pure-type game is generic (i.e., no two outcomes yield the same payoffs for any player), in order to avoid unnecessary complications that might obscure the main results. We begin by recalling here a classical result that we will be using throughout this article.

### Theorem

(Static equilibria and the replicator dynamics; Hofbauer and Sigmund, 1998; Webb, 2007). *Let* 𝔸 *be the set of asymptotically stable equilibria of the replicator dynamics (eq. 5)*, ℕ *the set of symmetric Nash equilibria of the game, and* 𝔽 *the set of equilibria (fixed points) o f t he replicator d ynamics (eq. 5). We then have*

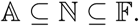

The above theorem’s purpose is mainly to allow us making statements about the replicator dynamics by using easier arguments based on static payoff comparisons. A simple proposition that we can now prove is the following one.

### Proposition 2

(Impossible invasion of costly plasticity). *In games where the replicator dynamics converge to an equilibrium point in pure strategy, costly plasticity cannot invade a population at the equilibrium of the pure-type game.*

*Proof.* Using the above theorem, we know that an asymptotically stable point of the pure-type game is necessarily a NE. Thus, when entering a monomorphic equilibrium population, a plastic player may either conform to the NE by playing the best-response *β*(*i*) = *i* to the resident *i*, or not conform to the NE by playing **z**_*i*_ *≠ β*(*i*). In the latter case player *p* will necessarily obtain a lower payoff than the NE payoff, *π*(**z**_*i*_, **x**_NE_) *< π*(**x**_NE_, **x**_NE_) which then implies that *w*_*p*_(**x**_NE_) *< w*_*i*_(**x**_NE_). If, in the former case, player *p* instead chooses the best-response *β*(*i*), its payoff will be the NE payoff *π*(**z**_*i*_, **x**_NE_) = *π*(**x**_NE_, **x**_NE_) but its fitness will still be lower than that of the resident because of the cost of plasticity, i.e., *π*(**x**_NE_, **x**_NE_) *-k < π*(**x**_NE_, **x**_NE_), so *w*_*p*_(**x**_NE_) *< w*_*i*_(**x**_NE_).

This result implies that a pure NE of the pure-type game is also a NE of the extended game with plastic types. While it is impossible for plastic types to invade pure NE, it is always possible to invade a mixed NE, as the next proposition shows.

### Proposition 3

(Coexistence of plasticity and pure types in games with mixed NE). *In games with no pure symmetric Nash equilibrium, there always exists a strategy* **z** *∈* (Δ𝒜)^*n*+1^ *such that plasticity persists in the population at equilibrium, i.e., there exists a 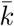 such that, for all 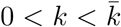*, lim_*t→∞*_ *φ*(*t*, **x**_0_)_*p*_ *>* 0, *for all* **x**_0_ *∈* int(Δ^*n*+1^).

*Proof.* Since the game has no pure NE, there is always, for each pair of actions *j ∈𝒜, i ∈𝒜*, an action *r*_*j*_ *∈𝒜* such that *π*(*i, j*) *≤ π*(*r*_*j*_, *j*). If *i* = *β*(*j*), then *r*_*j*_ = *i* = *β*(*j*), while if *i ≠ β*(*j*), then *r*_*j*_ can be any action that ensures *π*(*i, j*) *< π*(*r*_*j*_, *j*). If for each *j ∈𝒜*, the plastic type chooses to play **z**_*j*_ = *r*_*j*_ (by a slight abuse of notation), then in eq. 9 is satisfied, and plasticity can invade the mixed NE. This proves that any mixed NE can be invaded by a plastic type for sufficiently small plasticity costs. Since the game has no pure NE, the plastic type can also invade every monomorphic population of pure types, i.e., the equilibria **e**_*i*_ for all *i ∈𝒜* are locally unstable. This proves that all equilibria where plasticity is absent are locally unstable under the replicator dynamics, and that lim_*t→∞*_ *φ*(*t*, **x**_0_)_*p*_ *>* 0.

We just defined the conditions for plasticity to invade populations of pure types, but these do not guarantee local stability of plasticity. The following proposition establishes a necessary condition for the *inability* of plasticity to be immune against invasion by pure types.

### Proposition 4

(Impossible stability of plasticity). *If there exists i* ∈ *𝒜 such that plasticity cannot resist invasion by i, i.e. such that w*_*p*_(**e**_*p*_) *< w*_*i*_(**e**_*p*_), *then* (*i, i*) *is the symmetric Pareto-efficient equilibrium of the game.*

*Proof.* If plasticity cannot resist invasion by *i* ∈ 𝒜, this means that

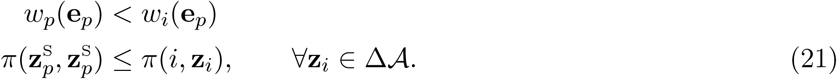

By way of contradiction, suppose that (*i, i*) is not the symmetric Pareto-efficient equilibrium of the game. Then there exists 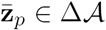 such that 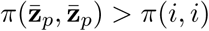. Hence a strategy of the plastic type with 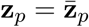 and **z**_*i*_ = *i* would guarantee local stability of plasticity. But, by eq. 21, this contradicts our assumption that plasticity is not locally stable against the invasion by *i*. This proves the result.

The converse is not true. If (*i, i*) is Pareto-efficient, it might still be possible to find s trategies o f the plastic type such that it resists invasion by *i*. Taking any of the three social dilemmas above, cooperation is Pareto-efficient, but it is always possible to find a s trategy t hat g uarantees l ocal s tability of plasticity.

### 6.1 Zero-sum games

From the discussion of the previous sections, it seems that if the two tasks of maximizing one’s own payoff and minimizing the other’s payoff are two compatible endeavours, plastic types can achieve global stability. Hence, the class of *strictly competitive games* would seem to favor plastic types. A game is strictly competitive if, for all *k ∈𝒜* and any pair of strategies *i ∈𝒜, j ∈𝒜*, the inequality *π*(*i, j*) *> π*(*k, j*) implies that *π*(*j, i*) *< π*(*j, k*). In particular, maximizing one’s own payoff is equivalent to minimizing the other’s payoff. Note that in a strictly competitive game, all strategy pairs are Pareto-efficient since switching from any action that induces a higher payoff for a focal player induces a loss for his opponent. It has been shown that strictly competitive games are affine transformations of zero-sum games (Adler et al., 2009). We next show that in the class of symmetric zero-sum games without pure NE, there always exists a strategy of the plastic type that guarantees its global stability under the replicator dynamics.

#### Proposition 5

(Global stability of costly plasticity in zero-sum games). *In zero-sum games with no pure symmetric NE, there always exists a strategy* **z** *∈* (Δ𝒜)^*n*+1^ *of the plastic type such that it is globally stable under the replicator dynamics.*

*Proof.* In order to show that plasticity is globally stable, we proceed by constructing a strategy that has the three properties required by Prop. 1:

1. It invades every monomorphic population of pure types.
2. It is locally stable against the invasion by pure types.
3. It invades a population at the mixed NEs of the pure type game.

To construct our winning plastic strategy, we begin by choosing the response of the plastic type against every pure type. Since the game has no pure symmetric equilibrium, there always exists, for each *i ∈𝒜*, at least one action *j*_*i*_ *∈𝒜, j*_*i*_ *≠ i*, such that *π*(*j*_*i*_, *i*) *>* 0. Let the response of our plastic type be to precisely choose such a *j*_*i*_, i.e. **z**_*i*_ = *j*_*i*_ (note that *j*_*i*_ need not be the best-response to *i*). To completely specify the strategy of the plastic type, we further need to define **z**_*p*_, the response of the plastic type to itself. Let us set **z**_*p*_ = **e**_*i*_, for arbitrary *i ∈𝒜*. We now verify that the three conditions for global stability are fulfilled by our plastic strategy.

As to condition (1), any pure type obtains a payoff as a resident of *π*(*i, i*) = 0, while our plastic type obtains *π*(**z**_*i*_, *i*) = *π*(*j*_*i*_, *i*) *>* 0. This means that *π*(**z**_*i*_, *i*) *> π*(*i, i*) for all *i ∈𝒜* and thus guarantees that plasticity invades any monomorphic population of pure types.

Regarding condition (2), the payoff of the plastic type as a resident is *π*(**z**_*p*_, **z**_*p*_) = *π*(*i, i*) = 0, for any *i ∈𝒜*, by definition of a symmetric zero-sum game. On the other hand any mutant pure type obtains *π*(*i*, **z**_*i*_) = *π*(*i, j*_*i*_) = *-π*(*j*_*i*_, *i*) *<* 0 against the plastic type. Hence *π*(**z**_*p*_, **z**_*p*_) *> π*(*i*, **z**_*i*_) for all *i ∈𝒜*, which guarantees local stability against the invasion by pure types.

Finally, condition (3) requires that the plastic type invades a population at the mixed NE. Since our plastic type obtains a positive payoff against each pure type because *π*(**z**_*i*_, *i*) = *π*(*j*_*i*_, *i*) *>* 0, for each *i ∈𝒜*, the expected payoff of a plastic type in any polymorphic population – including the population at NE frequencies – is also strictly positive, being a convex combination of positive payoffs. However the expected payoff of a pure type in a population where the frequencies of pure types correspond to the the mixed NE of the zero-sum game is *π*(**x**_NE_) = 0. Hence *w*_*p*_(**x**_NE_) *> w*_*i*_(**x**_NE_) for any *i* in support of the NE, which guarantees invasion of NE by plasticity. This completes the proof.

Note that the requirement that the game possesses no pure symmetric NE excludes symmetric 2 *×* 2 zero-sum games, since these games necessarily have a dominant action. Another remark is that the proof is also valid for strictly competitive games since our argument is independent of affine transformations to the payoffs of the game. This remark allows us to apply our results to the famous *good* Rock-Paper-Scissors game, where winning yields a payoff of *a*, while loosing induces a loss of *b*, and *a > b* (Sandholm, 2011), which entails the payoff matrix

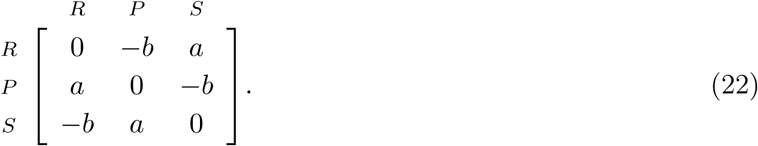

In such a game, the interior NE, **x**_NE_ = (1/3, 1/3, 1/3), is globally stable under the replicator dynamics, with solution trajectories displaying damped oscillations around this equilibrium. In Fig. 4, we show the effect of adding a plastic type with a strategy described in the proof of Prop. 5 to the good RPS game. In this game, this creates a plastic type that plays the best-response to any pure type and plays any *i ∈𝒜* against itself such that **z**_*i*_ = *β*(*i*) for any *i ∈* {*R, P*, *S*} and **z**_*p*_ = (1, 0, 0) [the choice of always playing *R* is arbitrary, in accordance with the proof of Prop. 5]. Even though the plastic type cannot invade on the faces of the simplex of dimension 2 (because symmetric 2 *×* 2 zero-sum games necessarily have a dominant action, and hence a pure symmetric NE), it achieves a larger payoff than any fully mixed population. This explains why it is globally stable in the interior of the simplex.

**Fig. 4:**
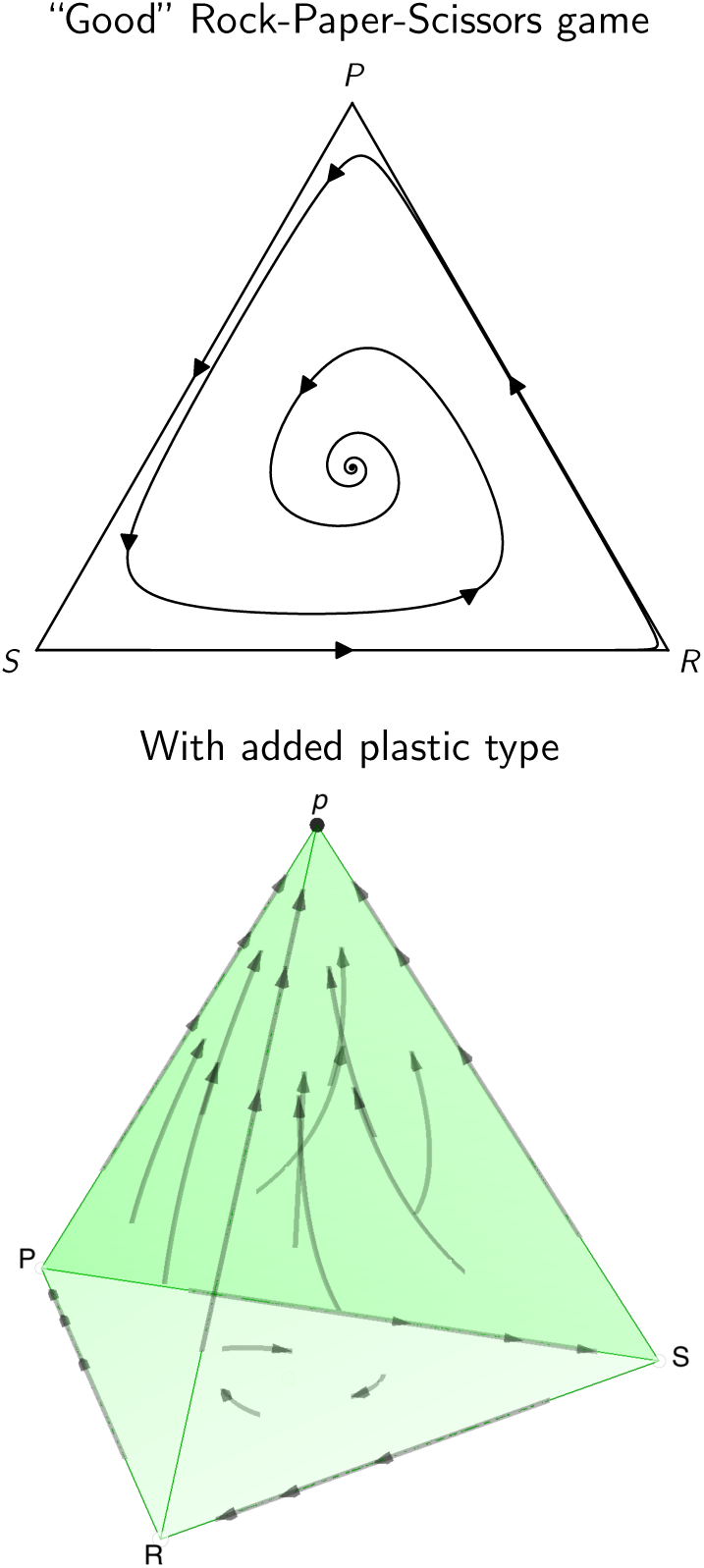
Replicator dynamics in the original good RPS (top), and with added plastic type (bottom). The 3-strategy simplex was produced using our own code (available), while the 4-strategy simplex was produced using Dynamo (Franchetti and Sandholm, 2013). Parameter values: Payoff matrix of eq. 22 with *a* = 2 and *b* = 1; **z**_*R*_ = (0, 1, 0); **z**_*P*_ = (0, 0, 1); **z**_*S*_ = (1, 0, 0); **z**_*p*_ = (1, 0, 0).

## 7 Discussion

In this work, we considered the introduction of plastic types in populations consisting of simple individuals adopting pure strategies in evolutionary symmetric games. We showed that despite having the capacity of responding in any possible way to other types in the population, there does not exist a general strategy for plasticity to eliminate pure types from the population. There are several explanations to this. The first one is that in games with pure symmetric Nash equilibria, costly plasticity cannot invade an equilibrium population. Moreover, we demonstrated that there is often a conflict between optimizing invasion success and optimizing stability for plastic types, the best illustration of this result being the Stag-Hunt game, where these two tasks (optimizing invasion ability versus stability) lead to totally different behavioral responses of the plastic types. We also saw that the success of plasticity is dependent on the class of games played in the population. There are games, such as zero-sum games without pure symmetric NE, where there always exists a strategy that grants plasticity global stability under the replicator dynamics. On the other hand, there are games, such as certain Mutualism games, where plasticity can neither invade an equilibrium population nor be stable against the invasion by pure types.

In our more detailed investigation of games of cooperation, we found that in the Prisoner’s Dilemma and the Snowdrift game, plastic types should cooperate with themselves with a strictly positive probability in order to be immune against the invasion by pure types. However, invasion success of pure populations by plasticity is generally uncorrelated to playing a cooperative strategy. In the Stag-hunt game it is just impossible for costly plasticity to invade an equilibrium population. However, costless plasticity is able to do so, and can even establish cooperation (the payoff-dominant outcome) in a population that was initially at equilibrium for defection (the inferior risk-dominant outcome). This result is similar to what Robson (1990) found in an earlier work (we compare our work with that of Robson, 1990, in more detail below).

It is insightful to look at our results through the lens provided by previous studies of the evolution of plasticity. In particular, previous work (e.g., Gomulkiewicz and Kirkpatrick, 1992) has revealed that variance in fitness is a key factor driving the evolution of plasticity. In our setting, we notice that in games with pure Nash equilibria (NE), the population does not experience variance in payoffs at equilibrium. Hence, it makes sense that there is no advantage to adapt behavior to different circumstances (in accordance with Prop. 2), since there is only one circumstance which is to face the pure type that is evolutionary stable. However, in games without pure NE, there is always variance in payoffs because one might be matched with any type in the support of the mixed NE of the game. This explains why plasticity can always invade a mixed NE, because it has the possibility to express different responses to the different existing types, while everyone else is forced to the NE payoff. Altogether, these results suggest that in general the ability to collect more fitness-relevant information is not always beneficial, as long as this information is costly to acquire, as other studies of the evolution of learning have previously shown (Wakano et al., 2004; Nakahashi et al., 2012; Aoki and Feldman, 2014; Dridi and Lehmann, 2015).

The proposition that individuals with more information may have an evolutionary advantage in social interactions is not new. Robson (1990) developed a model where his plastic type could only distinguish between plastic and non-plastic types and condition their action on this signal. He assumed that plasticity was costless and found in particular that plasticity can invade a population at the inferior equilibrium of a coordination game, and lead to the superior payoff-dominant equilibrium; however his plastic type could not co-exist with the superior pure type that plays the payoff-dominant action, in contrast to what we found. Banerjee and Weibull (1995) studied a very similar model to ours, but constrained the plastic type to play a best-response to every pure type. We saw that this best-response assumption is not the best strategy for plastic types when it comes to evolutionary stability. This phenomenon is exemplified by the Snowdrift game, where playing the best-response to every pure type does not allow plasticity to be locally stable against the invasion of pure mutants; but there are Snowdrift games where playing a strategy different than the best-response allows plasticity to be not only locally but globally stable. Our model is thus an extension of the ideas developed in these two earlier studies (Robson, 1990; Banerjee and Weibull, 1995).

From a broader perspective, given the recent renewed interest for the evolution of strategies in the repeated Prisoner’s dilemma (Stewart and Plotkin, 2016; Adami and Hintze, 2013; Press and Dyson, 2012; Stewart and Plotkin, 2013), our work helps bridge the gap between these recent studies and classical work on the evolution of pure non-plastic strategies in one-shot games (Hofbauer and Sigmund, 1998). Indeed, in order to be able to express a repeated-game strategy, an organism must first have the capacity to express a plastic social phenotype. Another recent trend is the focus on environmental variation in social evolution (Ashcroft et al., 2014; Dridi and Lehmann, 2014; Weitz et al., 2016; Estrela et al., 2018), and our study reminds that environmental variation is already embedded within social evolution through frequency-dependent payoffs (Graves and Weinreich, 2017), which is why we find that plasticity may evolve even in the absence of variation in the game payoffs.

Our model finally introduces new questions and calls for further research on the topic. Most notably, we ignored evolution of the plastic response itself, but it is natural to imagine that if one plastic type successfully invades, other plastic mutants with a different behavioral response might appear and displace the original plastic type; there is then no guarantee that the optimal plastic type will also be evolutionarily stable. Indeed, Robson (1990) showed that in the Prisoner’s dilemma, if a first plastic mutant might successfully invade and establish cooperation, the introduction of further plastic mutants might lead to the loss of cooperation. A possible avenue for future research would be to see if this result applies to our setting as well, in the Prisoner’s dilemma and other games. Another extension of our work would be to consider the possibility that the game payoffs change as a function of time, and in this setting a plastic type would be able to condition its action not only on the type of the opponent but also on the type of game being played. Heller (2004) modelled a situation where some individuals could detect the state of the environment and the opponent’s type in a context of a fluctuating game, but she assumed that plastic types (that she calls learning agents) only play the best-response to pure types. It would be interesting to see what would happen if we do not constrain the plastic response to be the best-response, since we saw in our model that the best-response does not always yield the best outcome for plasticity. These and other extensions will help us better understand the evolution of complex strategies in realistic changing environments.

## Acknowledgements

Support from ANR-Labex IAST is gratefully acknowledged.

